## Supplemental data for "Brain natriuretic peptide improves heart regeneration after infarction by stimulating cardiomyocyte renewal"

Short Title: BNP treatment increases cardiomyocyte cell number.

**Address for correspondance:**
* Nathalie Rosenblatt-Velin, PhD
CHUV, Division d'Angiologie
Département Cœur-Vaisseaux
Institut de Physiologie, Bugnon 7a,
1005 Lausanne, Switzerland


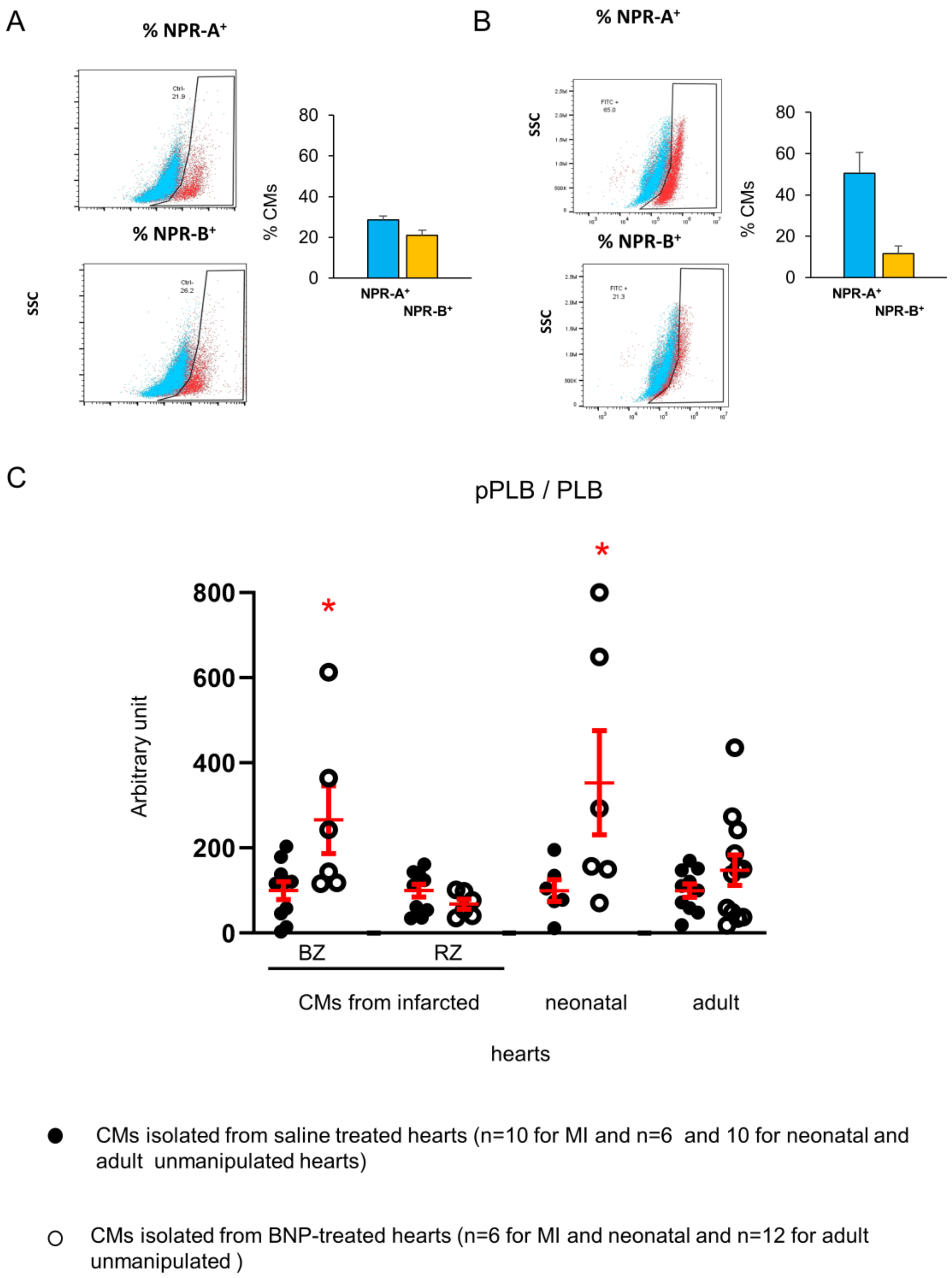


**<u>Supplemental Figure 1.</u> Neonatal and adult cardiomyocytes express NPR-A and NPR-B receptors and are able to respond directly to BNP stimulations. A-B.** NPR-A or NPR-B expression assessed by flow cytometry analysis on isolated neonatal (**A;** n=6 different isolations) and adult (**B;** n=4 different isolations) cardiomyocytes. Results represented as means ± SEM **C.** pPLB/PLB ratio in CMs isolated from adult infarcted, unmanipulated neonatal or adult hearts treated *in vivo* with saline (black circles) or BNP (white circles). Blots stained with antibodies against phospho phospholamban (pPLB), phospholamban (PBL) and Tubulin (used as loading control). Results of BNP treated CMs expressed relatively to the average of saline treated CMs. Individual isolations represented and the means ± SEM represented in red * p<0.05 versus CMs isolated from saline treated hearts.

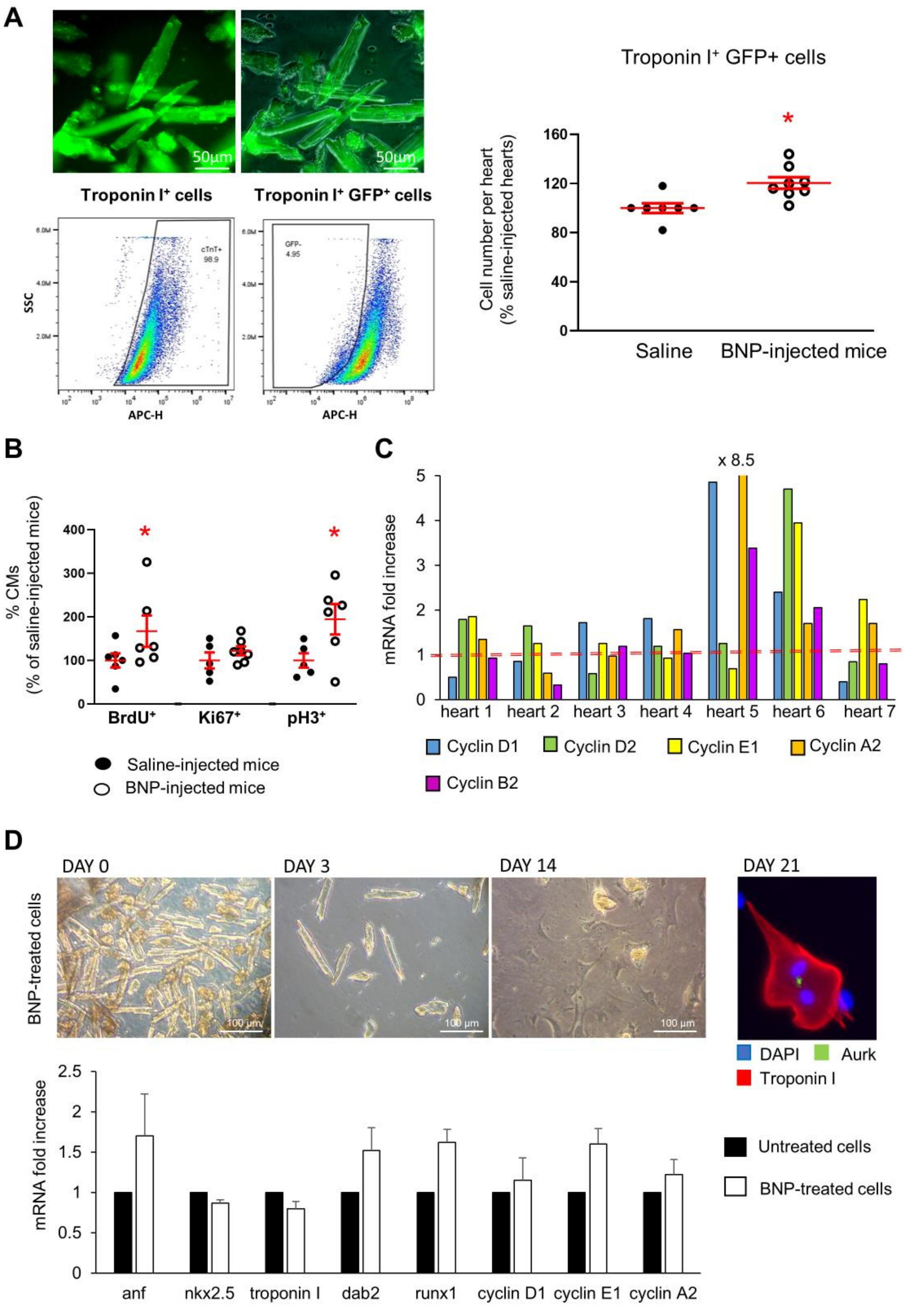


**<u>Supplementary Figure 2.</u> BNP treatment stimulates cardiomyocytes in unmanipulated adult hearts to re-enter in the cell cycle**. **A.** CMs isolated from unmanipulated Myh6 MerCreMer mice, treated 2 weeks with BNP or saline and injected with Tamoxifen 2 weeks before BNP injections. Flow cytometry analysis performed on these isolated CMs with an antibody against Troponin I. CMs identified as Troponin I$^+$ GFP$^+$ cells, counted**. B**. Immunostainings using antibodies against $\alpha$ actinin and BrdU or Ki67 or pH3 performed in adult unmanipulated hearts from saline or BNP-injected mice. The percentages of CMs expressing these proliferative markers measured on at least 10 pictures per mouse. **A-B:** Individual values represented and the means ± SEM represented in red. n=6 saline and BNP-injected hearts. **C**. mRNA expression coding for cyclin D1, D2, E1, A2 and B2. Results of CMs isolated from adult BNP-treated hearts expressed as fold increase above the levels in CMs isolated from saline-treated hearts (represented by the red dotted line, n= 8 saline-injected adult hearts). 7 different CM cell isolations from BNP-treated hearts represented**. D**. Adult cardiomyocytes isolated from 6-week-old unmanipulated C57BL/6 hearts cultured *in vitro* with or without BNP (10 nM). Representative pictures showing the evolution of the cell culture and the presence of Aurkb$^+$ Troponin I$^+$ cells in BNP-treated cells. CMs analyzed by qRT-PCR after 7 days of culture for the expressions of mRNAs coding for the atrial natriuretic peptide (anf), nkx2.5, dab2 and runx1, both re-expressed during CM dedifferentiation, and the different cyclins (D1, E1 and A2). n=3 different cell cultures. Results of BNP-treated CMs related to those of untreated cells. Data are means ±SEM. * $p < 0.05$ versus untreated cells.

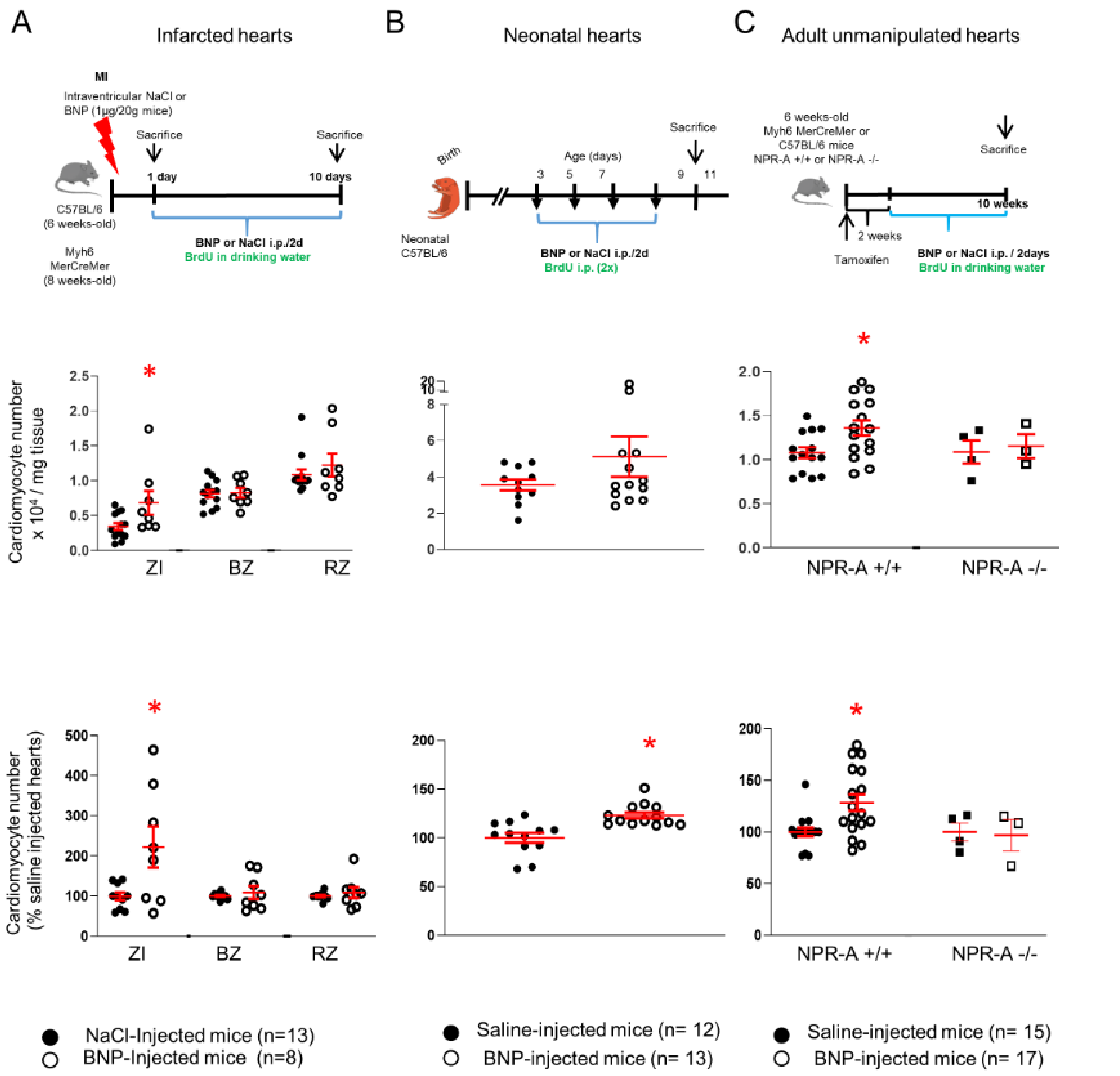


**<u>Supplemental Figure 3.</u> BNP injections in infarcted adult mice (A), in neonatal (B) and adult (C) unmanipulated mice lead to increased number of cardiomyocytes.** Cardiomyocytes isolated from BNP or saline injected heart counted and the number related to the tissue weight or to the number found in saline-injected mice. Individual isolation represented and the means ± SEM in red ***** p<0.05 versus saline treated hearts.

**Supplemental Figure 4. BNP treatment increases the number of neonatal cardiomyocytes *in vitro*. A.** Representative pictures of 2-weeks-old neonatal CM cell culture treated with different BNP concentrations (0-1000 nM). **B**. CM cell number related to the number of untreated CMs of the same isolation. n= 6 different cell cultures. For each experiment, the number of Troponin I+ cells / 0.9 $mm^2$ counted on at least 10 different pictures. **C.** Cardiomyocyte cross-sectional area measured in BNP treated cells and related to the area of untreated CMs. At least 60 CMs were measured per condition and cell culture (n= 6). **D.** Percentages of mononucleated CMs in untreated and BNP-treated conditions. At least 2700 CMs originating from 5 different cell cultures were evaluated in each condition. **E-F**. Detections by immunostainings of CMs (Troponin I+ cells) expressing Ki67, pH3 or Aurora B (Aurkb) proliferative markers in untreated or BNP-treated (10-100 nM) cultures. **G.** Percentages of CMs expressing these markers in BNP-treated cells related to untreated cells (n=6-8 different cell cultures). **H.** mRNA expression coding for cyclin genes D1, E1, A2 and B2. Results of BNP-treated CMs related to those of untreated cells.

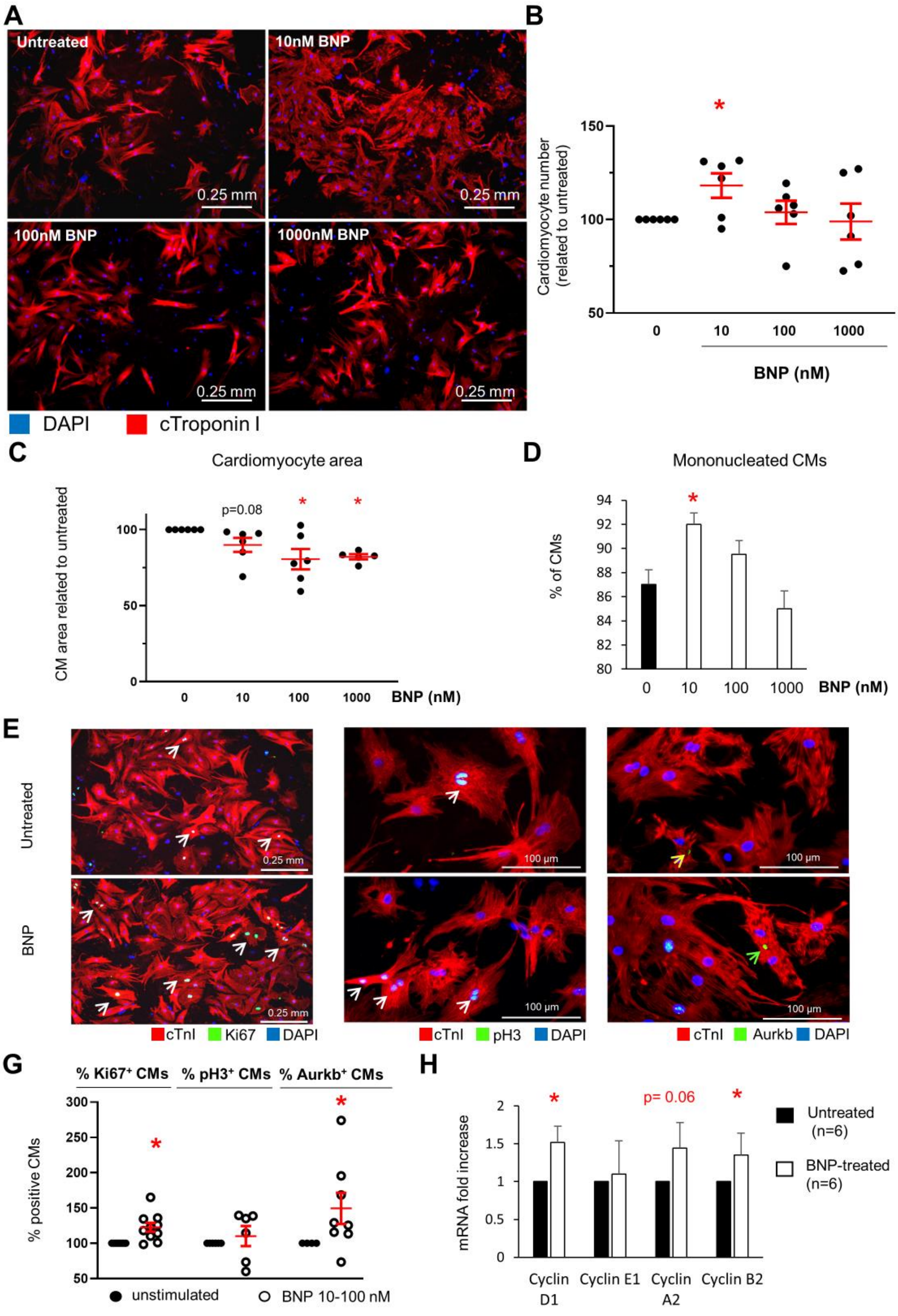

**Supplementary Figure 5**. Movie generated by time lapse microscopy showing the cytokinesis of a neonatal GFP positive cardiomyocyte stimulated with BNP (yellow arrow).

### Supplementary methods

#### Tamoxifen injections in adult Myh6 MerCreMer mice

Before surgery and/or BNP treatments, adult Myh6 MerCreMer mice (6-week-old) were injected with Tamoxifen (Sigma, T5648) at 40mg/kg to activate the Cre recombinase. Tamoxifen was dissolved in ethanol to a concentration of 100 mg/ml and emulsified in peanut oil to a final concentration of 10 mg/ml. 1mg Tamoxifen/25g body weight was injected intraperitoneally (i.p) to adult mice one time. Two weeks after tamoxifen injection, 90-95% of pre-existing CMs expressed the GFP protein (1).

#### Isolation of adult cardiomyocytes

Rapidly after sacrifice, the chest of the mice was opened, descending aorta and inferior vena cava were cut, the aorta was clamped and EDTA buffer was perfused into the right ventricle. Thereafter, the heart was removed from the chest and the left ventricle was perfused with enzymatic solution containing collagenase II (120U/ml), collagenase IV (120 U/ml) (Worthington, Biochemical Corporation, USA) and protease XIV (0.05mg/ml or 0.175U/ml) (Sigma). In each experiment, a full digestion of the heart was obtained. After digestion, cardiac cells were dissociated by gentle pipetting, filtered and collected after 25 min sedimentation. For infarcted hearts, the tissue was separated into the three zones (ZI, BZ and RZ). The three different area were weighted before gentle pipetting.

Total CM cell numbers were counted before sedimentation by two different people and in blinded experiments. The CMs were easily identified based on their size. Round and rod shaped CMs were counted.

After 25 min sedimentation, CMs were used for molecular analysis, western blot analysis, flow cytometry analysis or fixed with 2.5% PFA 15 min at room temperature (RT) and stained with DAPI to evaluate the frequency of mono and bi-nucleated CMs. Flow cytometry analysis based on Troponin I staining demonstrated that NMCs contamination in the CM fraction after sedimentation corresponds to ≤5% of the total cells and that sedimentation enriched the CMs fraction into rod-shaped CMs (i.e 90% of rod-shaped CMs were collected at the end of the sedimentation).

#### Neonatal cardiomyocyte culture

Neonatal (1-2 days) C57BL/6 pups were sacrificed and CMs were isolated by enzymatic digestion according to the method previously described (2, 3). Briefly, the chest was opened and the heart removed. After separation of the atria from the rest of the heart, the ventricles were minced and digested with 0.45mg/ml collagenase (Worthington, Biochemical Corporation, USA) and 1 mg/ml pancreatin (Sigma). After 3 rounds of digestion, cells were plated 2 times for 45 min in order to separate CMs (non adherent cells) from the non-myocyte cells (NMCs). Thereafter, CMs were plated on gelatine (0.1%) coated plates and cultured in 3:1 mixture of DMEM and Medium 199 (Invitrogen Corp, San Diego, CA, USA) supplemented with 10% horse serum (Oxoid), 5% fetal bovine serum (Invitrogen), 10 mM Hepes, 100U/ml penicillin G. To homogenize the experiment, 70000 CMs were plated per well.

Immediately after plating, neonatal CMs were treated or not with BNP for 14 days. To inhibit NMC proliferation and to work on 95% pure CMs, cells were exposed the first 7 days of culture to cytosine-β-

D-arabinofuranoside (AraC, 1 mg/ml). After 7 days of culture, AraC was removed until 14 days of culture. CMs were cultured at 3% $O_2$ in a standard incubator or in a hypoxia chamber (Stem Cell Technologies) flushed with 3% $O_2$/5% $C0_2$/92% $N_2$ (Carbagas). Medium was replaced 1-2 times/week.
CMs were treated with 3 different concentrations of BNP: 10nM, 100nM and 1000nM and compared to untreated cells. Quantitative RT-qPCR, western blot analysis as well as immunohistochemistry studies were performed after 14 days of culture.

**Flow cytometry analysis**

Adult CMs were isolated using enzymatic digestion (see Materiel and Methods). Adult CMs isolated from Myh6 MerCreMer hearts were fixed with 5.5% formaldehyde and permeabilized with 0.5% saponin. CMs were stained during 20 min at RT with anti-troponin I antibody (Supplemental Table 1). Cells were analyzed with CytoFLEX (Beckmann Coulter) cytometer and data generated using FlowJo 10 software. The number of $cTnI^+$ $GFP^+$ cells per heart was obtained by relating the percentage of the $cTnI^+$ $GFP^+$ CMs acquired by flow cytometry analysis to the total number of CMs in the heart. Cells were stained with DAPI and only the DAPI negative cells (living cells) were analysed.

**Immunohistochemistry**

Cells or OCT heart sections (5µm-thick cryosections) were washed in PBS 1X and fixed in paraformaldehyde (2%) or formol (4%) for 10 min at RT. After 10 min of permeabilization (0.3% Triton x-100 in PBS) and one hour of blocking with normal serum, sections were probed with primary antibodies overnight at 4°C (Supplemental Table 1). The second day, the secondary antibody was added on cells or heart sections. Slides were mounted with Dabco (Sigma) and pictures were captured with Nikon Eclipse TS100 or 90i microscope. Images were processed with Adobe Photoshop CC2015. For the detection of BrdU incorporation, heart slides were fixed 10 min in 2% PFA, DNA was denatured 1h at RT in HCl 2N before neutralisation in Na Borate 0.1M pH=8.5 during 2x5 min. Then, sections were probed with primary antibodies overnight at 4°C (Supplemental Table 1).

**Time lapse microscopy**

1 day after birth, Myh6 MerCreMer/Tomato-EGFP pups were injected intraperitoneally with tamoxifen (1mg / 2g). 1 day after, CMs were isolated and were plated on laminin (10µg/ml) substrate. CMs were cultured in media containing 3:1 mixture of DMEM and Medium 199 (Invitrogen Corp, San Diego, CA, USA), supplemented with 10% horse serum (Oxoid), 5% fetal bovine serum (Invitrogen), 10mM Hepes and, 100U/ml penicillin G at 20% O2. After isolation, CMs were stimulated with BNP (100 nM). 48 hours post plating, CMs were transferred to an OkoLab environment system with temperature and $CO_2$ stage incubator. CMs expressing EGFP were targeted and images were taken each 40 min for 48 hours using the Nikon Eclipse Ti2 inverted microscope and NIS-Elements software (Nikon).

**qRT-PCR**

Total RNA was isolated from heart tissue, CM cell cultures and isolated CMs using TRI-Reagent (Zymo Research). cDNA was synthesized from RNA using PrimeScript RT reagent kit (Takara Bio Inc). Polymerase chain reactions (PCR) were performed using the SYBR Premix Ex Taq polymerase (Takara

Bio Inc) with the ViiA$^{TM}$7 Instrument (Applied Biosystems). Results were obtained after 40 cycles of a thermal step protocol consisting of an initial denaturation 95°C (1s), followed by 60°C (20s) of elongation (α-skeletal actin has an elongation time of 30s at 60°C). The sequences of primers were reported in Supplemental Table 2. All results were normalized with the 18S housekeeping gene (Δ CT values). Means of ΔΔ CT ($\Delta\ CT_{BNP}$ - $\Delta\ CT_{saline}$) values (versus untreated cells or NaCl treated mice) were calculated and results were represented as $2^{-\Delta\Delta CT}$. Statistics were performed on ΔΔ CT individual values. SEM fold increase was calculated using $2^{-\Delta\Delta\ CT\ high\ values}$- $2^{-\ means\ of\ \Delta\Delta\ CT}$ (4).

**Supplemental Table 1 :** Antibodies used in flow cytometry analysis, immunohistology and western blot analysis.

| | Species | Dilution | Reference | Technics |
|---|---|---|---|---|
| **Primary antibodies** | | | | |
| α-actinin | mouse | 1/50 | Sigma A7811 | Immunohistology |
| Aurkb | rabbit | 1/1000 | Abcam ab139188 | Immunohistology |
| BrdU | rat | 1/100 | Abcam ab6326 | Immunohistology |
| cleaved caspase 3 | rabbit | 1/400 | Cell Signaling 9661 | Immunohistology |
| GFP | rabbit | 1/1000 | Abcam ab290 | Immunohistology |
| Phospho Histone H3 (S10) | rabbit | 1/100 | Millipore 06-570 | Immunohistology |
| Ki67 | rat | 1/1000 | ebioscience 14-5698-80 | Immunohistology |
| Laminin | rabbit | 1/200 | Sigma L9393 | Immunohistology |
| NPR-A | rabbit | 1/50 | Abcam ab70848 | Immunohistology |
| NPR-B | rabbit | 1/100 | Abcam ab139188 | Immunohistology |
| Troponin I | goat | 1/100 | Santa Cruz Biotechnology SC-8118 | Immunohistology |
| Akt | rabbit | 1/1000 | Cell Signaling | Western Blot |
| Phospho-Akt | rabbit | 1/500 | Cell Signaling | Western Blot |
| Bax | rabbit | 1/1000 | Cell Signaling | Western Blot |
| Bcl-2 | rabbit | 1/1000 | Cell Signaling | Western Blot |
| cleaved caspase 8 | rabbit | 1/1000 | Cell Signaling | Western Blot |
| cleaved caspase 3 | rabbit | 1/1000 | Cell Signaling | Western Blot |
| Erk | rabbit | 1/3000 | Cell Signaling | Western Blot |
| NPR-B | goat | 1/20 | Santa Cruz SC-34421 | Flow cytometry |
| NPR-A | rabbit | 1/50 | Abcam ab70848 | Flow cytometry |
| NPR-C | mouse | 1/100 | GeneTex | Flow cytometry |
| Phospho-Erk | rabbit | 1/2000 | Cell Signaling | Western Blot |
| p38 | rabbit | 1/1000 | Cell Signaling | Western Blot |
| Phospho-P38 | rabbit | 1/500 | Cell Signaling | Western Blot |
| phospholamban | mouse | 1/1000 | Abcam | Western Blot |
| Phospho-phospholamban | rabbit | 1/500 | Millipore | Western Blot |
| Troponin I | goat | 1/50 | Abcam ab56357 | Flow cytometry |
| Tubulin | mouse | 1/10000 | Sigma T5168 | Western Blot |
| **Secondary antibodies** | | | | |
| Anti-rabbit Alexa 488 | donkey | 1/1000 | Molecular Probes A21206 | Immunohistology |
| Anti-rabbit Alexa 594 | donkey | 1/1000 | Molecular Probes A21207 | immunohistology |
| Anti-goat Alexa 594 | donkey | 1/1000 | Molecular Probes A11058 | Immunohistology |
| Anti-mouse Alexa 647 | goat | 1/1000 | Molecular Probes A21240 | Immunohistology |
| Anti-rat 647 | donkey | 1/500 | Jackson Immuno 712-605-150 | Immunohistology |
| Anti-rat Alexa 488 | donkey | 1/1000 | Molecular Probes A21208 | Immunohistology |
| Anti-rat biotinylated | goat | 1/200 | Vector BA-9400 | Immunohistology |
| Steptavidine Alexa 594 | | 1/1000 | Molecular Probes S11227 | Immunohistology |
| Anti-rabbit Alexa 680 | goat | 1/5000 | Molecular Probes A21109 | Western Blot |
| Anti-mouse IRDye 800 | goat | 1/10000 | Rockland Immuno-chemicals 610-132-121 | Western Blot |
| Anti-goat APC-conjugated | chicken | 1/10 | R&D systems F0108 | Flow cytometry |
| Anti-mouse IgG2b FITC | goat | 1/1000 | Molecular Probe A21141 | Flow cytometry |

**Supplemental Table 2:** List of primers used in qRT-PCR.

| Gene | Forward primer | Reverse primer | Product size (bp) |
|---|---|---|---|
| **anf** | ACAGGATTGGAGCCCAGAGC | GTCCATGGTGCTGAAGTTTATTC | 337 |
| **acta1** | TGGACTTCGAGAATGAGATGG | TCGTCCTGAGGAGAGAGAGC | 509 |
| **cyclin A2** | ATGTCAACCCCGAAAAACTG | GCAGTGACATGCTCATCGTT | 157 |
| **cyclin B2** | AGCTCCCAAGGATCGTCCTC | TGTCCTCGTTATCTATGTCCTCG | 116 |
| **cyclin E1** | GAAAGAAGAAGGTGGCTCCGAC | GTTAGGGGTGGGGATGAAAGAG | 190 |
| **cyclin D1** | TGAGAACAAGCAGACCATCC | TGAACTTCACATCTGTGGCA | 71 |
| **cyclin D2** | GGATGATGAAGTGAACACACTCAC | GGATCTTCCACAGACTTGGATCC | 180 |
| **dab2** | TGCTCGTGATGTGACAGACA | AGGGTCATTAGGGCCTCACT | 225 |
| **hif1α** | CTGTCATCTCACTATGGGCA | CCAAGTCCGAGCAGGAATTT | 259 |
| **myh6** | AACCAGAGTTTGAGTGACAGAATG | ACTCCGTGCGGATGTCAA | 130 |
| **myh7** | ATGAGACGGTGGTGGGTTT | CTTTCTTTGCCTTGCCTTTG | 117 |
| **nkx2.5** | CAAGTGCTCTCCTGCTTTCC | GTCCAGCTCCACTGCCTTCT | 130 |
| **npr1** | CCAATTATGGCTCCCTGCT | CGGTACAAGCTCCCACAAAT | 198 |
| **npr2** | TCATGACAGCCCATGGGAAA | GGTGACAATGCAGATGTTGG | 209 |
| **npr3** | GGCTCAATGAGGAGGATTACGTG | AATCTTCCCGCAGCTCTCGATG | 555 |
| **runx1** | GATGGCACTCTGGTCACCG | GCCGCTCGGAAAAGGACA | 298 |
| **troponin T** | GCGGAAGAGTGGGAAGAGACA | CCACAGCTCCTTGGCCTTCT | 127 |
| **18S** | ACTTTTGGGGCCTTCGTGTC | GCCCAGAGACTCATTTCTTCTTG | 96 |
